# Temperature-dependent modulation of odor-dependent behavior in three drosophilid fly species of differing thermal preference

**DOI:** 10.1101/2023.04.11.536399

**Authors:** Steve B. S. Baleba, Venkatesh Pal Mahadevan, Markus Knaden, Bill S Hansson

## Abstract

Rapid and ongoing climate change increases global temperature and impacts both feeding and reproduction in insects. The sense of smell plays an important underlying role in these behaviors in most insect species. Here, we aimed to investigate how changing temperatures affect odor detection and ensuing behavior in three drosophilid flies: *Drosophila novamexicana*, *D. virilis* and *D. ezoana*, species that have adapted to life in desert, global and subarctic climates, respectively. Using a series of thermal preference assays, we confirmed that the three species indeed exhibit distinct temperature preferences. Next, using single sensillum recording technique, we classified olfactory sensory neurons (OSNs) present in basiconic sensilla on the antenna of the three species and thereby identified ligands for each OSN type. In a series of trap assays we proceeded to establish the behavioral valence of the best ligands and chose guaiacol, methyl salicylate and isopropyl benzoate as representatives of a repellent, attractant and neutral odor. Next, we assessed the behavioral valence of these three odors in all three species across a thermal range (10-35 C), with flies reared at 18°C and 25°C. We found that both developmental and experimental temperatures affected the behavioral performance of the flies. Our study thus reveals temperature-dependent changes in odor-guided behavior in drosophilid flies.

## Introduction

The olfactory system of insects underpins a large number of evolutionarily critical behaviours such as search for food, oviposition substrates, shelters, and mates, and avoidance of harmful organisms including parasitoids, predators and pathogens ^1^. Olfactory detection is carried out with an array of olfactory sensory neurons (OSNs) located on the antennae ^2^ and maxillary palps ^3^. In case of *Drosophila melanogaster*, the third antennal segment, the funiculus, is covered with a large number of hair-like structures, sensilla, which house dendrites of OSNs (Nava Gonzales et al., 2021). These OSNs express distinct olfactory receptors (ORs), which dimerize with the co-receptor Orco ^4,5^. While OSNs housed in trichoid sensilla respond predominantly to pheromones ^6^, and those associated with coeloconic sensilla mainly to amines and carboxylic acid ^7^, general food and oviposition substrate odour detection mainly takes place in OSNs present in basiconic sensilla.

A significant consequence of human activities can be observed in terms of climate change and consistently increasing global temperatures, which in turn might affect odour guided behavior of insects. Insects are ectotherms, meaning that their biology is closely related to environmental temperatures ^8^. Any changes in the external temperature can thus affect the physiology of the olfactory sensory system leading to modulation of behaviour triggered by the system ^9^. For instance, at the behavioural level, the sensitivity of *D. melanogaster* to ethanol increases or decreases when flies are acclimatized to heat or cold, respectively ^10^. Furthermore, electrophysiological recordings revealed changes in odour detection at the OSN level ^11^. Another study showed that exposing *D. melanogaster* flies to 30°C for 24 hours altered the expression level of genes involved in odour detection, causing a reduction in avoidance of high ethanol concentrations ^12^. Conclusions drawn from these studies were based on flies reared at 21°C, acclimatized at 15°C (cold treatment) or 30°C (heat treatment) and subsequently tested at 24°C using electrophysiology and behavoural studies. Similar studies conducted in other insect orders are based on insect acclimatization as well. Linn et al.^13^ reared the moths *Grapholita molesta* (Lepidoptera: Tortricidae) and *Pectinophora gossypiella* (Lepidoptera: Gelechiidae) at 25°C or 26-27°C, after which they were tested in a wind tunnel at 20°C and 26°C. These two temperatures induced differential specificity in male pheromone-directed responses. Processing of olfactory stimuli has also been shown to be temperature-dependent in other insects ^14^.

The effect of developmental temperature and/or experimental temperature on odour detection and odor-dependent behavior in drosophilid species evolutionarily adapted to different climates has so far received little attention. In the present study, we used three closely related species belonging to the *virilis* group; *Drosophila ezoana*, *D. novamexicana* and *D. virilis*, which have adapted to live in subarctic, desert and global climates, respectively. Despite their different climatic preferences, all three species feed and breed on slime flux, saps and decaying bark of tree species belonging preferably to the Salicaceae and Betulaceae families ^15^.

We first assessed the temperature preference of the three species when reared at 20, 23 and 25°C. Next, we used a panel of 57 ecologically relevant odorants to characterize response profiles of distinct OSN classes and thereby identified active ligands for each OSN class. Further, we established the behavioral valence of key ligands identified for each OSN type using a two choice trap assay. Lastly, we chose three odors that elicited attraction, aversion or neutrality, respectively, across the three species and tested whether behavioral responses would be affected by developmental and/or experimental temperatures.

## Materials and methods

### Flies stocks and husbandry

We obtained *Drosophila ezoana* (stock number: E-15701) from the fly stocks of Kyorin university, Japan (https://shigen.nig.ac.jp/fly/kyorin/index_ja.html), while *D. novamexicana* (stock number: 15010-1031.08) and *D. virilis* (stock number: 15010-1051.00) were obtained from the National *Drosophila* Species Stock Center of Cornell University (https://www.drosophilaspecies.com/). We reared *D. ezoana* at 20°C, 16h Light: 8h Dark, *D. virilis* at 23°C 12h Light: 12h Dark, and *D. novamexicana* at 25°C 12h Light: 12h Dark. All flies were fed on autoclaved cornmeal-yeast-sucrose-agar food.

### Temperature preference assay

An apparatus with thermally conductive material that can be heated in one end and cooled at the other offers the most suitable way of measuring thermal preference in animals ^16^. Based on this, we designed a thermal gradient choice bioassay based on a design developed by Lynch et al.^17^. The apparatus (Supplementary figure 1 A, right) was composed of 10 aluminium channels measuring each 24 × 1 × 0.8 cm and separated from each other by 0.5 cm (Supplementary figure A, bottom) and two heat and cool Quick-Ohm generators (https://www.quick-ohm.de/) to allow the establishment of a thermal gradient (going from 10 to 40°C) along each aluminium channel. On top of the aluminium channels, a sheet of plexiglass was placed to maintain flies in the channels whilst enabling continuous observations. The plexiglass was drilled in the middle with 10 holes of 5 mm in diameter (covered with a screw) to ease the introduction of individual flies into each channel. To avoid phototaxis effects during the experiment, we kept the apparatus in darkness using a darkened cage, and infrared light and camera (Supplementary figure 1 A, right and left).

**Figure 1.**
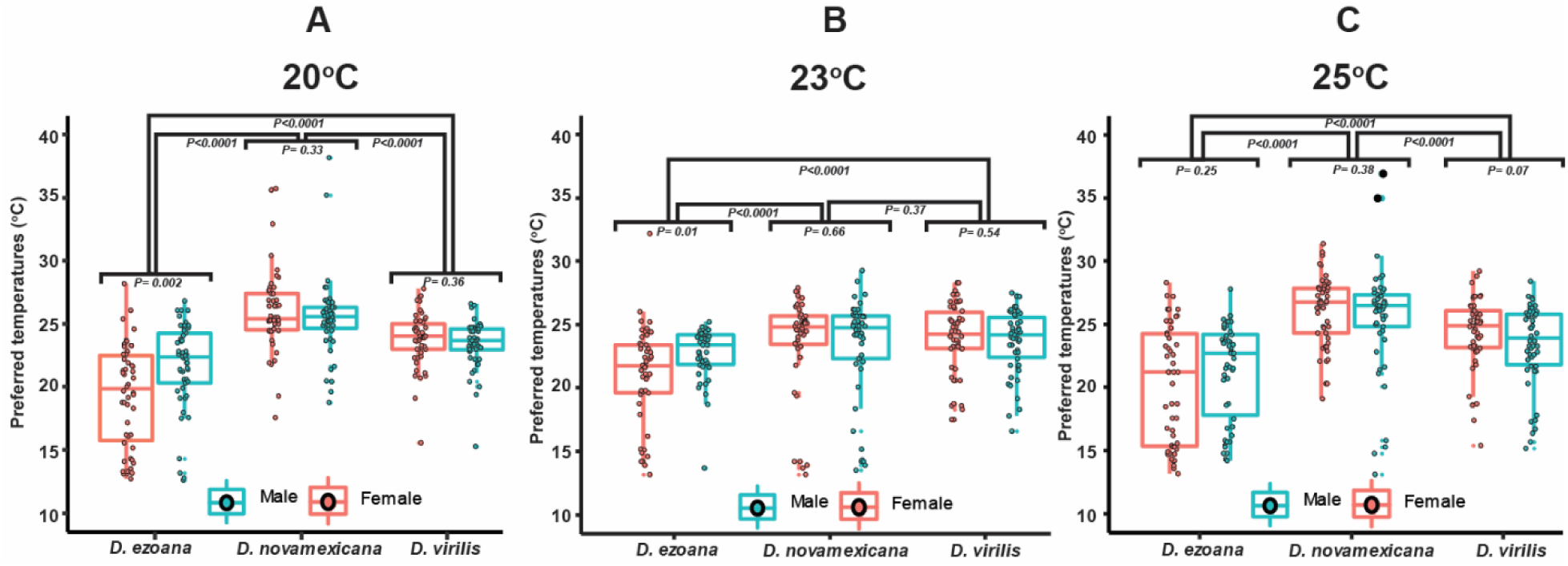
Temperature preference of the three *Drosophila* species. Boxplot illustrating the preferred temperatures of males and females of *D. ezoana* **(A)**, *D. novamexicana* **(B)** and *D. virilis* **(C)** when reared at 20 °C (left), 23°C (middle) and 25 °C (right). Boxplot whiskers indicate ± 1.5 interquartile range limits. Dots in each boxplot indicate individual temperature data points (n=50 individuals). Significance (*P<0.05*) was tested in an unpaired t-test (comparison between sexes of each species) and a one-way ANOVA followed by SNK multiple comparison post hoc tests (comparison across species).

The movements of the three species were first observed without turning on the heat and cold generators. To do this, we used 50 males and 50 females (4-to-6 days old) of each species and gently introduced them individually (using a mouth aspirator) inside each channel (Supplementary figure 1 A, bottom). Their movements along the channels were recorded for 30 minutes using a camera placed on top of the bioassay. At the end of each recording, the channels were cleaned with ethanol before introducing the next set of 10 individuals. Using the recorded videos, the movement of the flies was tracked by determining their position in the channels every two minutes. To check for the consistency of the thermal gradient in the assay, we recorded (using a data logger thermometer equipped with two probes (https://www.tcdirect.de/)) the temperature of 12 fixed positions in each channel 1, 2 and 3 hours after turning on the heat and cold generators.

After these control experiments, the heat and cold generators were again turned on. After 10 minutes a thermal gradient, spanning from 10 to 40°C was established along the aluminium channels. Individual flies were then introduced into each channel. Fifty males and 50 females of each fly species were tested and their movements were video taped for 30 minutes. We used the temperature of the region of the channel where an individual fly spent most of its time as a proxy for its preferred temperature ^17,18^. To determine possible effects of developmental temperature, *D*. *ezoana*, *D. novamexicana* and *D. virilis* individuals used in this experiment were reared at three different temperatures; 20, 23 and 25°C.

### Electrophysiology

Single sensillum recordings were performed following the protocol described by Olsson and Hansson^19^ . Briefly, adult flies of each species were restrained in a 100µl plastic pipette tip with the wide end closed with dental wax and the narrow end cut to allow only the head with antennae to protrude. The preparation was fixed on dental wax placed on a microscope slide with the ventral side of the fly facing upward. The funiculus of the antenna was fixed with a sharpened glass capillary (placed between the second and third antennal segment) held with dental wax onto a cover slide, which in turn was held in place by dental wax on the microscope slide. Afterwards, the preparation was placed under a light microscope (BX51WI, Olympus, Tokyo, Japan) equipped with a ×50 magnification objective (LMPLFLN 50X, Olympus) and 4x eyepieces. The preparation was continuously flushed by clean air through a plastic tube of 0.4 cm diameter delivering a 1.5 l min^-1^ flux of charcoal-filtered and humified air. The tube ended approximately 6cm m from the preparation.

Stimulus cartridges were prepared by placing a circular filter paper (1.2 cm diameter) in the large opening end of a Pasteur pipette and pipetting 10 µl of odorant solution onto the paper before closing the pipette with a 1 ml plastic pipette closed at its small opening with wax to prevent evaporation. The antenna of the fly was stimulated by a 500 ms air pulse (0.6 l min^-1^) through the stimulation cartridge into the permanent air flux (1.5 l min^-1^).

To record action potentials electrochemically sharpened tungsten electrodes were used. The electrodes were sharpened using saturated potassium nitrite (KNO2) solution. The reference electrode was inserted into the eye of the fly with the aid of a manually controlled micromanipulator. With the aid of a motor-controlled DC-3K micromanipulator (Märzhäuser, Wetzlar, Germany) equipped with a PM-10 piezo translator (Märzhäuser), the recording electrode was inserted at the base of a sensillum. The electrical signal was amplified using a USB-IDAC connection to a computer (Syntech, The Netherlands). The frequency of action potentials during a 1-second pre- and post-stimulation period was established using Auto Spike software (Syntech, version 3.7). Approximately 200 sensilla from the funiculus of each of the three *Drosophila* species were recorded from, using the *D. melanogaster* sensilla distribution map as a reference Lin and Potter^20^. 2,6-dimethoxy-phenol, 3,4,5-trimethoxyphenol, oleamide, acetovanillone, salicylaldehyde and hexadecanamide were diluted in dimethyl sulfoxide (DMSO) at 10^-4^ concentration (1:10000 volume:volume), while all remaining odorants were diluted in mineral oil at 10^-4^ concentration. All synthetic odorants used were of the highest purity available from Sigma (www.sigmaaldrich.com) and Bedoukian (www.bedoukian.com). The entire odorant panel was tested on maximally 3 sensilla per fly.

### Behavioural assay

The single sensillum recordings allowed us to identify key odorants that elicited strong responses in each of the basiconic OSN types of the three *Drosophila* species. With this information as a base we turned to a dual choice bioassay to establish the valence (i.e. attraction, aversion, or no response) that the key odorants triggered in the three species. Trap assay experiments were performed in a climate chamber (23° C, 70% humidity, 12h Light: 12h Dark). Transparent plastic boxes of 500 ml (with 30 ventilation holes in the lid) held treatment and control traps made from small, transparent plastic vials (25 ml) with a paper cone (cut at the apex (4mm)), inserted into the opening of the vial and fixed with transparent tape. Treatment and control traps were loaded with a lid of an Eppendorf tube containing either 2µl of the odorant to be tested (diluted in 200 µl mineral oil at 10^-2^ concentration) or 200µl of solvent (mineral oil). For each species, we transferred 20 flies (10 males and 10 females, 4–6 days old, starved for 24 hours before the experiment) inside the large box. We counted the number of flies that had entered the traps and those that remained outside after 24 hours. With this data, an attraction index (AI) could be calculated as AI= (O-C)/20, where O is the number of flies that entered the trap containing the odorant and C is the number of flies that entered the trap containing the solvent. The AI ranged from -1 (maximum avoidance) to 1 (maximum attraction). Zero thus denoted no choice. For each species, 20 odorants were tested and for each odorant the trap assay was replicated 10 times.

Among the 20 odorants tested in the trap assay, we identified some that triggered attraction, some aversion, while some were neutral across all the three species. Next, we aimed to see whether the developmental and/or the experimental temperatures could modulate these behaviors. For this, the three species were reared at 18 and 25°C, after which the emerged adults (4–6 days old) were tested in trap assays as described above, with the particularity that all the experiments were conducted inside a thermo controlled Percival incubator (www.percival-scientific.com). The experiments were run at 10, 15, 20, 25, 30 and 35°C, all with 70% humidity and 12h Light:12h Dark. As stimuli, methyl salicylate (positive), guaiacol (negative) and isopropyl benzoate (neutral) were used. The compounds were diluted in mineral oil 10^-2^ concentration. No significant difference in release rate across the different temperatures and times was observed (Supplementary figure 4 and 5).

**Figure 4.**
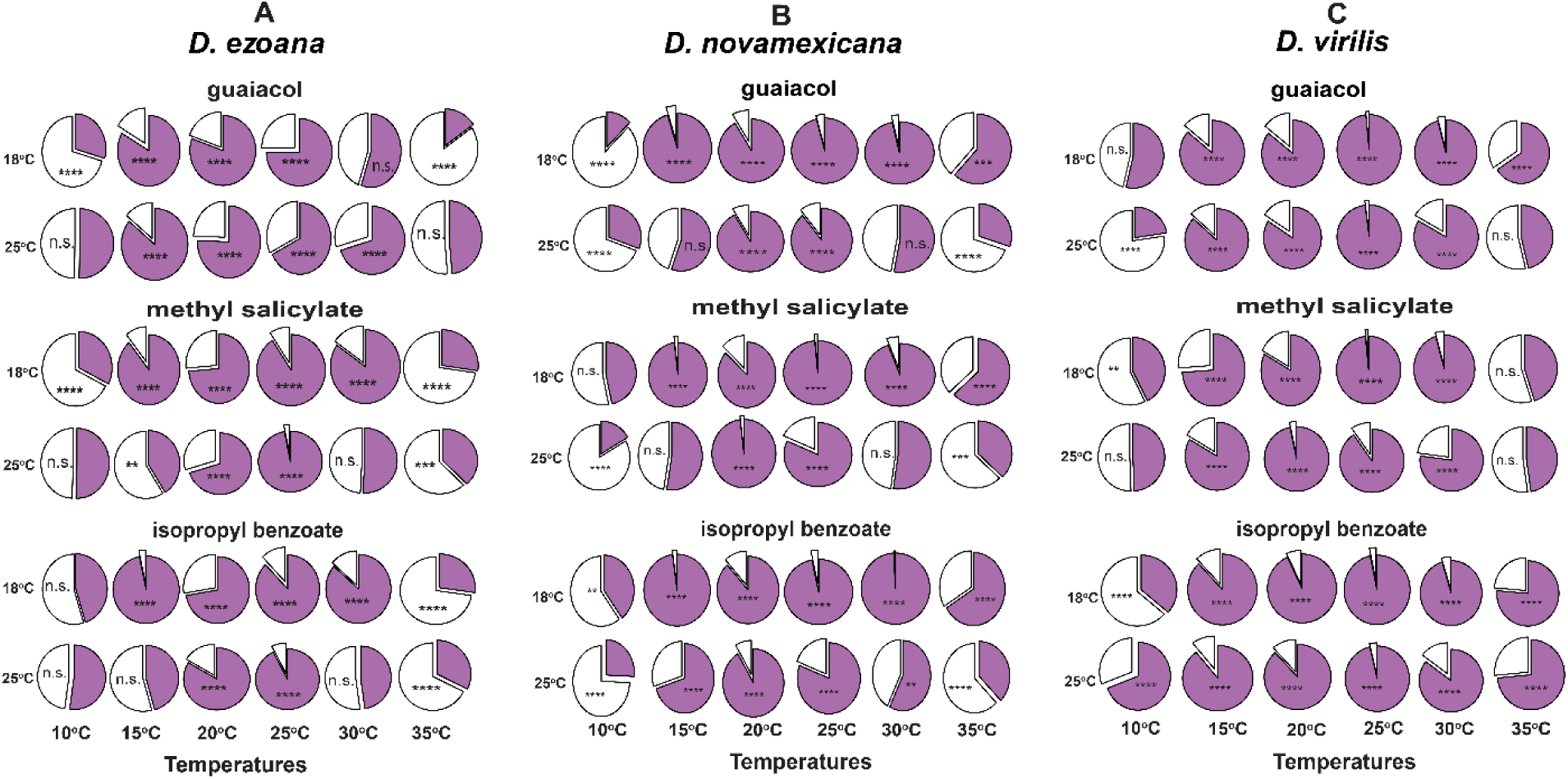
Effect of temperature on valence. Pie charts showing the proportion of *D. ezoana* **(A)**, *D. novamexicana* **(B)** and *D. virilis* **(C)** individuals found in the control and treatment container (purple portion) in comparison to the proportion of individuals found outside the containers (white portion). Proportions compared using the Chi-square test. The symbols n.s indicate non-significant differences; *, **, ***, **** depicts *P<0.05*, *P<0.01*, *P<0.001* and *P<0.0001,* respectively.

**Figure 5.**
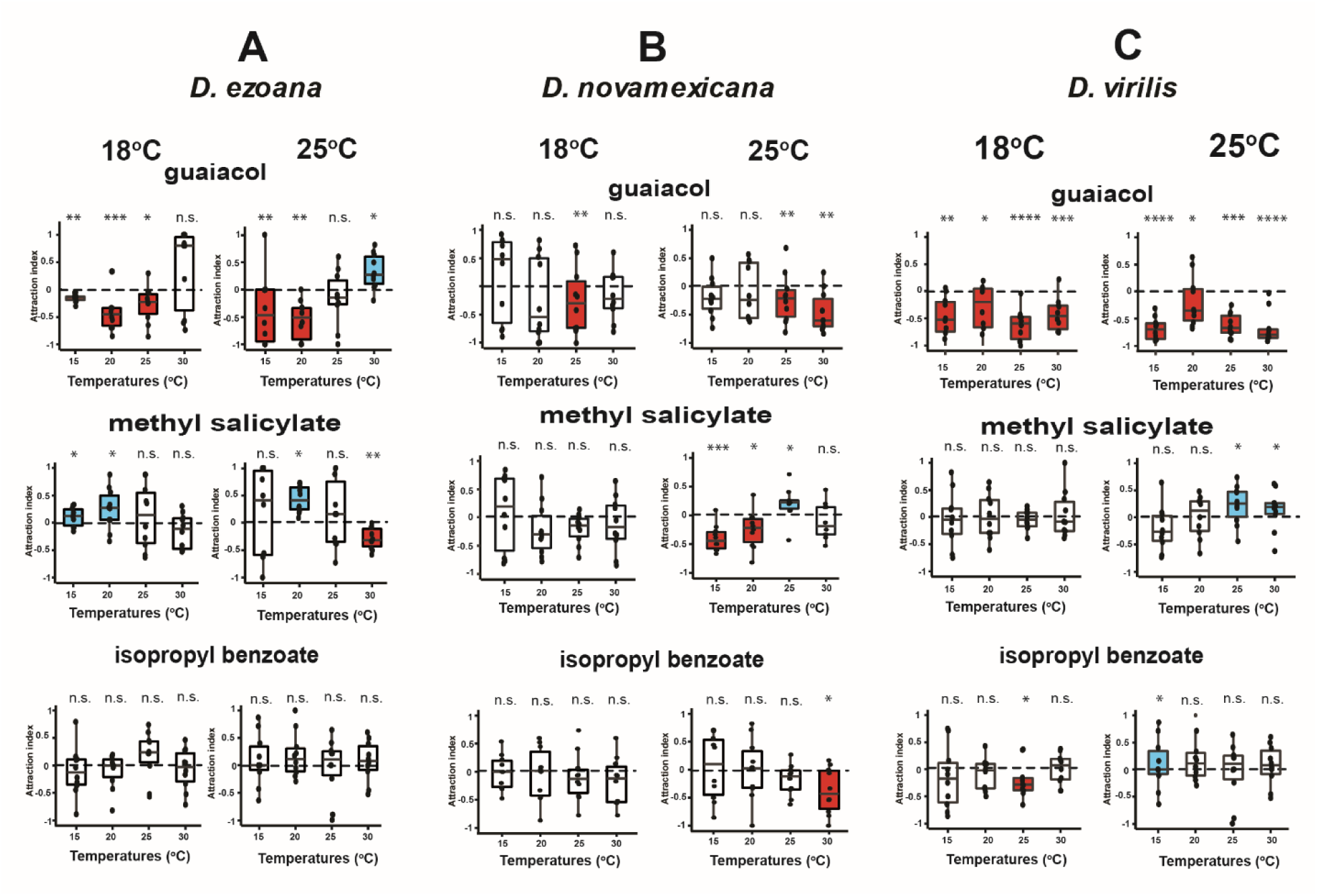
Effect of temperature on odorant valence. (**A**) Boxplot showing the temperature effect on aversion, attraction and neutral activity of guaiacol, methyl salicylate and isopropyl benzoate, respectively in *D. ezoana* reared at 18^°^C (left) and 25^°^C (right). (**B**) The same representation for *D. novamexicana*. (**C**) The same representation for *D. virilis*. The edges of the boxes are the first and third quartiles, thick lines mark the medians, and whiskers represent the data range. The dots in each box show individual data points (n=10 replicates, 20 individuals per replicate). Box plots coloured in red, blue and white depict aversion, attraction and no response, respectively. Significance of the attraction index to the theoretical mean 0 was tested using the one-sample t-test (normally distributed data: Shapiro test: *P>0.05*) or one sample Wilcoxon test (non-normally distributed data: Shapiro test: *P<0.05*).

### Data analysis

All statistical analyses were performed in R version 4.0.3 (R Core Team, 2020) and PAST version 3.09^21^ software and all graphs were assembled in Adobe Illustrator CC 2017 (version 21.0). The temperature preference data were normally distributed (Shapiro-Wilk test: *P> 0.05*) and their variances were homogeneous (Barlett test: *P>0.05*), therefore we ran an unpaired t-test to see whether males and females of the same species significantly preferred different temperatures. Analysis of variance was computed followed by Student-Neuman-Keuls (SNK) post hoc multiple comparisons tests using the R software package called ‘Agricolae’ (de Mendiburu, 2020) to compare the preferred temperatures across the three *Drosophila* species. With the SSR data recorded from each species, we employed the *heatmap ()* function embedded in the R software to generate the heatmaps illustrating the neuronal responses of each OSN type when individually stimulated by the 57 odorant panels. We used the one-sample t-test (normally distributed data: Shapiro test: *P>0.05*) or one-sample Wilcoxon (non-normally distributed data: Shapiro test: *P<0.05*) statistical tests to compare the attraction indices (calculated from the trap assay data) with the theoretical mean zero (0). Statistical results were considered significant when *P<0.05*.

## Results

### Temperature preference of Drosophila ezoana, D. novamexicana and D. virilis

The temperature preference of the three species bred at 20, 23 or 25°C temperature regimes was tested in mini thermal channels as described above. In initial control experiments, individual flies were released inside the channels in absence of a thermal gradient. After tracking their movements for 30 minutes, we observed that individuals of each species were distributed evenly inside the channels (Supplementary figure 1 B). When the heat and cold generators were engaged, a linear temperature gradient from 10 to 40°C was established inside each channel. The gradient was stable and remained unchanged even after 3h (Supplementary figure 1 C). When released inside the channels in presence of the thermal gradient, *D. ezoana* flies avoided the hot end of the gradient (Supplementary figure 2 A); whereas *D. novamexicana* avoided the cold end (Supplementary figure 2 B). *D. virilis* occupied an intermediate position in the gradient (Supplementary figure 2 C). The temperature experienced by the three *Drosophila* species during their preimaginal development did not influence their thermal preference at the adult stage (Fig. 1A, B, C). Males and females in general preferred the same temperature regime. Female *D. ezoana* did, however, show a significantly higher cold-loving propensity in comparison to the males of the same species. The three species investigated thus revealed clear temperature preferences matching their natural habitats.

**Figure 2.**
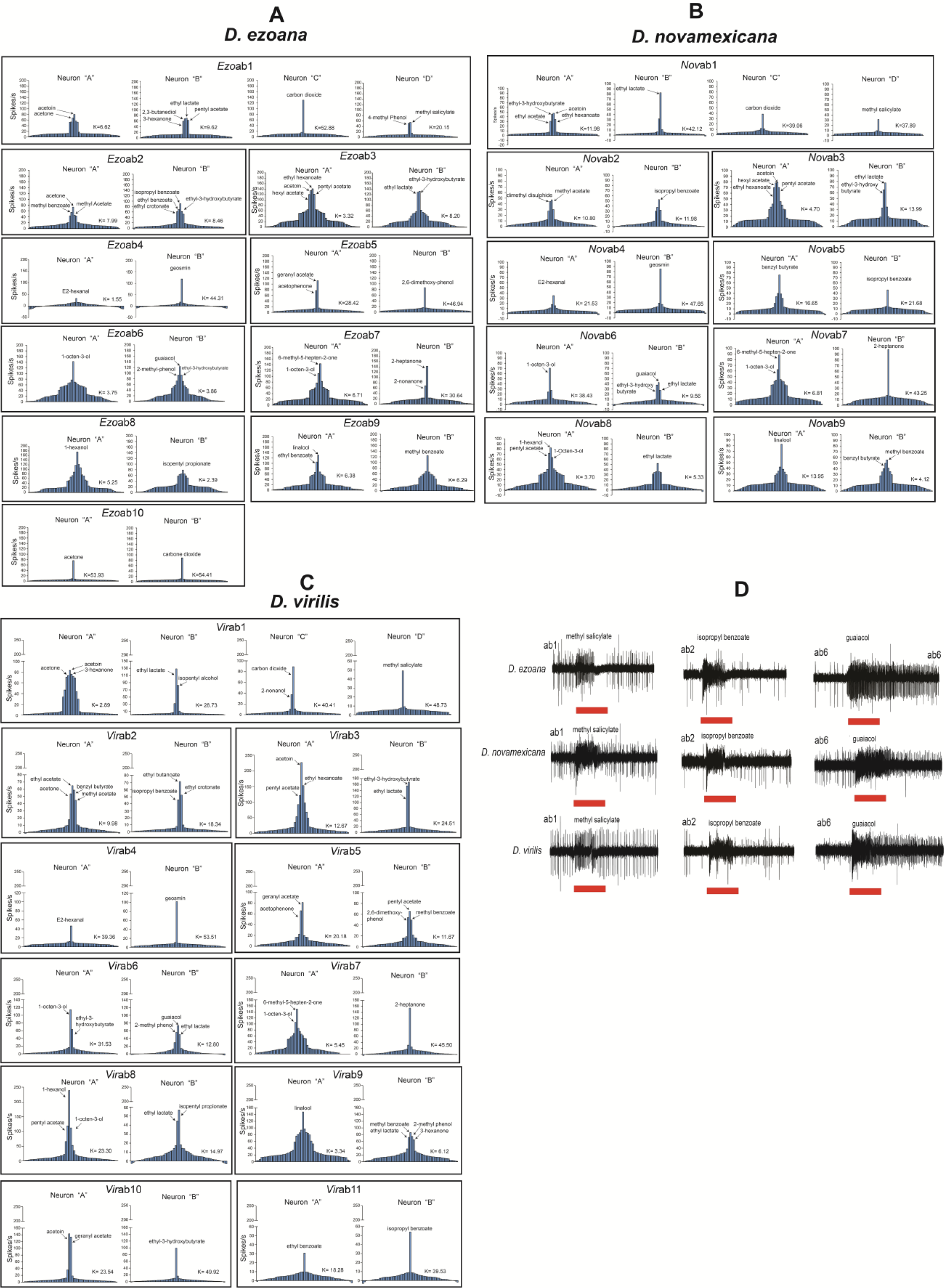
Tuning histograms of each olfactory sensory neuron present in basiconic sensillum types identified in *D. ezoana* (**A**), *D. novamexicana* (**B**) and *D. virilis* (**C**). x-axis represents the 57 compounds tested with the most active compound in the center; y-axis represents the number of action potentials (spikes) elicited.The kurtosis (k) value represents the ‘peakedness’ of each distribution curve. This value is high in narrowly tuned neurons, and low is broadly tuned neurons.**(D)** Representative single-sensillum recording (SSR) traces in *D. ezoana*, *D. novamexicana* and *D. virilis*, showing responses of ab1, ab2 and ab6 to methyl salicylate, isopropyl benzoate and guaiacol respectively.

### Physiological characterization of olfactory sensory neurons in antennal sensilla basiconica of the three species under study

We used the well-characterized olfactory sensory neuron (OSN) types associated with basiconic sensilla in *Drosophila melanogaster* as a vantage point for our dissection of the same type of neurons in *D. ezoana, D. novamexicana* and *D. virilis.* In comparison to *D. melanogaster*, where 10 basiconic types have been physiologically classified, we identified 10, 9 and 11 types, respectively in *D. ezoana*, *D*. *novamexicana*, and *D. virilis* (Supplementary Figure 3). When sensilla in these species contained an OSN collection with identical or almost identical response spectra as found in *D. melanogaster*, the sensillum types were given the same number as in *D. melanogaster* preceeded by a species identifier (*Ezo*, *Nov*, *Vir*). In a few cases, the key ligands of one OSN in a sensillum were identical to those of one *D. melanogaster* OSN, while the second OSN did not match the *D. melanogaster* pattern. Also, we observed that OSNs of *D. novamexicana* (Fig. 2 B) in general produced fewer action potentials as compared to *D. ezoana* (Fig. 2 A) and *D. virilis* (Fig. 2 C).

As in *D. melanogaster*, three large basiconic (LB) types were identified in *D. ezoana* (*Ezo*ab1, *Ezo*ab2, *Ezo*ab3), *D. novamexicana* (*Nov*ab1, *Nov*ab2, *Nov*ab3) and *D. virilis* (*Vir*ab1, *Vir*ab2, *Vir*ab3). In the three species, the *Ezo*ab1, *Nov*ab1 and *Vir*ab1 sensillum type housed the typical four OSNs, the “A” neuron, characterized by large action potentials or spikes, responding strongly to acetoin and acetone, “B”, “C” and “D” neurons responding respectively to ethyl lactate, carbon dioxide and methyl salicylate. In *D. ezoana*, a similar LB sensillum (*Ezo*ab10) was found. This sensillum type, however, contained only two responding neurons (A and B) strongly activated by acetone and carbon dioxide, respectively (Fig. 2A). This type was not found in *D. novamexicana* (Fig. 2B) or *D. virilis* (Fig. 2C). The *Ezo*ab2, *Nov*ab2 and *Vir*ab2 sensillum types responded to similar stimulus spectra. Their “A” and “B” OSNs responded strongly to methyl acetate and isopropyl benzoate, respectively. In addition, in *D. ezoana* and *D. virilis* acetone stimulated the “A” neuron, while in *D. novamexicana* it elicited a very weak response. In this species, the “A” neuron responded strongly also to dimethyl disulfide. The *Ezo*ab3, *Nov*ab3 and *Vir*ab3 sensillum types of all the species also exhibited similar response patterns. The “A” OSN responded strongly to ethyl hexanoate, acetoin and pentyl acetate, while the“B” OSN responded to ethyl lactate and ethyl-3-hydroxybutyrate.

Small basiconic (SB) sensillum types were present in all three *Drosophila* species. The *Ezo*ab4, *Nov*ab4 and *Vir*ab4 sensillum types housed two neurons “A” and “B” tuned to E2-hexanal and geosmin, respectively. The *Ezo*ab5 and *Vir*ab5 sensilla housed two OSNs: “A” that responded to geranyl acetate and acetophenone and “B” that responded to 2,6-dimethoxy-phenol. No similar sensillum type was found in *D. novamexicana*. Nonetheless, we found in this species a sensillum type that we called *Nov*ab5. It housed two neurons “A” and “B” responding respectively to benzyl butyrate and isopropyl benzoate. The *Ezo*ab6, *Nov*ab6 and *Vir*ab6 sensillum types housed two neurons “A” and “B” tuned to 1-octen-3-ol and guaiacol, respectively. The *Ezo*ab7, *Nov*ab7 and *Vir*ab7 hosted two neurons “A” and “B” that mainly responded to 6-methyl-5-hepten-2-one and 2-heptanone respectively. The *Ezo*ab8, *Nov*ab8 and *Vir*ab8 sensillum types responded strongly to 1-hexanol (A); while the B OSN responded to ethyl lactate and isopentyl propionate. The *Ezo*ab9, *Nov*ab9 and *Vir*ab9 sensillum types hosted an “A” neuron responding to linalool and a “B” neuron responding to methyl benzoate and a variety of other odorants. Lastly, in *D. virilis*, we found two sensilla: *Vir*ab10 and *Vir*ab11 absent in *D. ezoana* and *D. novamexicana*. The *Vir*ab10 sensillum housed two neurons; “A” tuned to acetoin and geranyl acetate, “B” tuned to ethyl-3-hydroxybutyrate. The *Vir*ab11 sensillum type displayed an “A” neuron tuned to ethyl benzoate and a “B” neuron primarily responded to isopropyl benzoate.

It must be noted that sensilla classification and its corresponding nomenclature is unique to individual species and must not be compared with the canonical *D. melanogaster* sensilla nomenclature and also within the three species. However, we found some OSN types conserved across the three species that are comparable to *D. melanogaster*. For instance, ab1 like class (diagnostic ligand: CO2), ab4 like (diagnostic ligand: geosmin), ab6 like (diagnostic ligand: guaiacol). However, these classes also demonstrate significant changes in terms of their co-innervating OSN types. For example, we observe that the A neuron in Dvirab1 like sensillum class responded strongest to acetoin as against ethyl acetate for the ab1A OSN type reported in D. melanogaster.

### Behavioral valence of the best ligands identified and its modulation by temperature

Using the best ligands identified for each class of OSN in the SSR recordings, we wanted to test their behavioural significance in the three species studied. Screening of the 20 best ligands in trap assays revealed that the ligand spectrum contained odors eliciting attraction, repulsion or neutral responses in *D. ezoana* (Fig. 3A), *D. novamexicana* (Fig. 3B) and *D. virilis* (Fig. 3C). Several compounds triggered different responses in the three species. For instance, 6-methyl-5-hepten-2-one and methyl benzoate elicited no response in *D. ezoana*, while in *D. novamexicana* and *D. virilis* they triggered repulsion. Acetoin and ethyl-3-hydroxybutyrate attracted *D. virilis*, while in *D. ezoana* and *D. novamexicana*, these compounds elicited no response. 1-hexanol attracted *D. novamexicana*, but in *D. ezoana* and *D. novamexicana* it triggered no response. Also, some compounds exhibited the same valence across the three species. Benzyl butyrate, linalool and methyl salicylate elicited attraction; guaiacol and hexyl acetate triggered aversion, while isopropyl benzoate and pentyl acetate did not induce any choice. No compounds elicited opposite responses.

**Figure 3.**
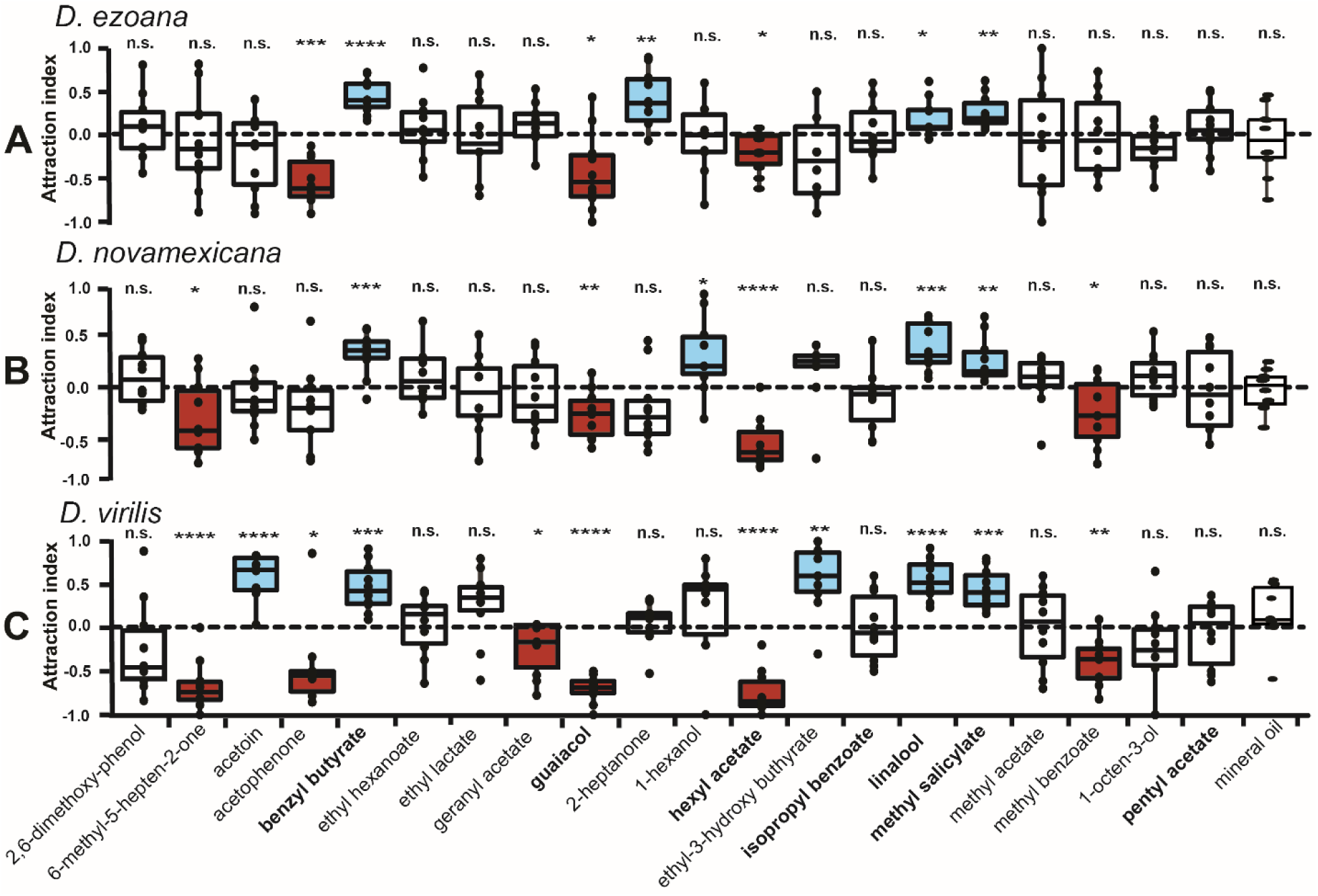
Valence value of key ligands of olfactory sensory neurons in *D. ezoana* (**A**), *D. novamexicana* (**B**) and *D. virilis* (**C**). Box plots show the attraction index (based on a two-choice trap assay (odour vs solvent) with **1** depicting maximum attraction, **0** neutrality and **-1** maximum repellency of the compound tested. In each boxplot, the ends of boxplot whiskers represent the minimum and maximum values of all the data and dots show individual data points (n=10 replicates, 20 individuals per replicate). Boxplots filled with blue, red and white colours depict respectively a attraction, repulsion and neutrality. Based on distribution of the data, one sample t-test (normally distributed data: Shapiro test: *P>0.05*) or one sample Wilcoxon (non-normally distributed data: Shapiro test: *P<0.05*) statistical tests were used to test the significance of the attraction index. The n.s. symbol denotes a non-significant difference of the attraction index to the theoretical mean 0; while *, **, ***, **** indicate statistical significance of the attraction index to the theoretical mean 0, with *P<0.05*, *P<0.01*, *P<0.001*, *P<0.0001* respectively.

We next asked whether the valence of these key ligands would change as a function of rearing and/or experimental temperatures. To address this, we conducted trap assays at 10, 15, 20, 25, 30, and 35°C using flies reared at 18 and 25°C. As representative attractant, repellent and neutral odorants, we used guaiacol (negative), methyl salicylate (positive) and isopropyl benzoate (neutral). In most of the cases, *D. ezoana* (Fig. 4A), *D. novamexicana* (Fig. 4B) and *D. virilis* (Fig. 4C) entered either into the control or treatment containers (the purple portion of the pie-chart) in trap assays conducted at 15, 20, 25 and 30°C. In trap assays performed at 10 and 35°C, we found a large proportion of flies outside the control and treatment containers (white portion of the pie-chart); suggesting that these two extreme temperatures considerably impaired the movement of the three species towards the trap containers. Based on this result we decided to exclude the extreme temperatures from further analysis.

When proceeding with a more detailed analysis of the behavioral responses in the two-choice assay, we found that in *D. ezoana* (Fig. 5A) reared at 18°C the repellency for guaiacol was significantly reduced (flies reacted neutrally) in trap assays conducted at 30°C. Among individuals reared at 25°C, the repellency disappeared in trap assays conducted at 25 and 30°C. At 30°C the flies even became attracted to guaiacol. High testing temperatures thus clearly affected the behavioral responses to guaiacol. For methyl salicylate we found conserved attraction in trap assays conducted at 15 and 20°C with *D. ezoana* reared at 18°C. However, when tested at 25 and 30°C, the flies exhibited neutral behaviour toward this compound. With individuals reared at 25°C, we only observed attraction to methyl salicylate in trap assays performed at 20°C, while at at 15 and 25°C these flies responded neutrally, and at 30°C they were even significantly repelled by this compound. The positive response to methyl salicylate thus had a very narrow temperature range. The neutral response to isopropyl benzoate in *D. ezoana* remained across the different developmental and experimental temperatures.

In *D. novamexicana* (Fig. 5B), flies reared at 18°C exhibited avoidance behaviour towards guaiacol when tested at 25°C. At 15, 20 and 30°C they responded neutrally. With individuals reared at 25°C, guaiacol was repellent at 25 and 30°C. At 15 and 20°C, the flies exhibited neutral behaviour. The negative esponse to guaiacol was thus abolished at lower temperatures. *D. novamexicana* reared at 18°C responded neutrally to methyl salicylate at all test temperatures. When reared at 25°C, we noted attraction to this compound only when tested at 25°C. These flies were instead repelled by methyl salicylate at 15 and 20°C, while responding neutrally at 30°C. As in *D. ezoana*, the positive response to methyl salicylate was extremely sensitive to the testing temperature. The neutral response to isopropyl benzoate did not change with test temperature when we used *D. novamexicana* reared at 18°C. However, when reared at 25°C, they showed repulsion at 30°C testing temperature.

In *D. virilis* (Fig. 5C), flies reared at 18 and 25°C avoided guaiacol under all different experimental temperatures. In the case of methyl salicylate, no attraction to this compound was observed with *D. virilis* developed at 18°C. Instead, they exhibited neutral behaviour across the different testing temperatures. When the flies were reared at 25°C, they were attracted to methyl salicylate at 25 and 30°C. At 15 and 20°C, they did not respond to the compound. The neutral response to isopropyl benzoate persisted across most experimental temperatures. However, *D. virilis* flies developed at 18°C showed aversion to this compound at 25°C and flies reared at 25°C were attracted at 15°C.

## Discussion

We aimed to study how insects can cope with the global increase in overall temperature from an olfactory point of view. We studied odor-dependent behavior in three *Drosophila* species representing arctic, desert and cosmopolitan distributions, namely *Drososphila ezoana*, *D. novamexicana* and *D. virilis*. After establishing differences in thermal preference range between the species, we used single sensillum recording technique to classify different basiconic sensillum types present on the antennae and identified best ligands for the corresponding olfactory sensory neurons (OSNs). In a series of trap assay experiments, we then established the behavior (attraction, aversion or no response) that these key ligands triggered in the three species. Finally, we tested both the effect of rearing and testing temperatures on the behavioral reactions by conducting trap assays at different testing temperatures using flies reared at 18 and 25°C and odorants previously identified as an attractant, repellent or neutral in the three species.

As expected from their natural habitats, *D. ezoana* showed a significant preference towards cooler temperatures, *D. novamexicana* towards warmer, while *D. virilis* preferred an intermediate temperature range. These thermal preference patterns remained consistent across flies reared at 20, 23 and 25°C, showing that despite different conditions of development, the thermal preference behaviour stayed constant. Maclean et al.^22^ stated that fundamental species-specific ecological characteristics such as temperature tolerance do not eclipse during laboratory maintenance. *D. ezoana*, *D. novamexicana* and *D. virilis* have adapted to life in the arctic, desert and global climates, respectively. In comparison, Rajpurohit and Schmidt ^23^ demonstrated that *D. melanogaster* from temperate areas exhibited a greater preference for cooler temperatures, while those from tropical habitats preferred higher temperatures. Similar patterns were found in other temperature-related parameters. *D. ananassae* flies living at high latitudes recovered faster from chill coma and more slowly to heat knockdown as compared to flies living at low latitudes ^24^. Obviously, *Drosophila* species in general show clear physiological climatic limits and geographical variation of genes involved in temperature preference and adaptation ^25,26^, that lead to a thermal preference matching the habitat they have evolved in.

As a base for the behavioral studies we performed an extensive screening of antennal sensilla basiconica. The sampling (200 in each species) yielded 10, 9 and 11 basiconic sensillum types in *D. ezoana*, *D. novamexicana* and *D. virilis*, respectively. The recordings revealed both similarities and differences from the 10 basiconic types physiologically classified in *D. melanogaster* ^4,27–29^*, D. mojavensis* ^30^ *and D. suzukii* ^31^. The differences observed likely stem from the fact that *Drosophila* flies colonise environments and display ecologies characterised by different types of sensory information that have shaped the molecular, physiological and anatomical organization of their sense of smell. Despite our extensive recordings, sampling effects might also play a role.

In general, we observed both gain and loss of some sensillum types in the three *Drosophila* species studied here. In *D. ezoana*, we identified a large sensillum type (*Ezo*ab10) containing two neurons “A” and “B” tuned to acetone and carbon dioxide, respectively, and thus seemingly lacking two of the OSNs present in the “normal” ab1 type. Across all three species, the *Ezo*ab3, *Nov*ab3 and *Vir*ab3 sensilla showed some functional deviation from ab3 sensilla of *D. melanogaster*. In *D. melanogaster*, the “A” and “B” OSNs of this sensillum type respond mainly to ethyl hexanoate and 2-heptanone, respectively ^27,32^. In our study, we found these two neurons in two distinct sensillum types. In the first sensillum type (*Ezo*ab3, *Nov*ab3 and *Vir*ab3), the ethyl hexanoate-responding “A” neuron innervated the same sensillum lymph as a “B” neuron excited by ethyl lactate and ethyl-3-hydroxybutyrate. In the second sensillum type (*Ezo*ab7, *Nov*ab7 and *Vir*ab7), the 2-heptanone-responding “B” neuron was housed in the same lymph as an “A” neuron responding mostly to 6-methyl-5-hepten-2-one and 1-octen-3-ol. In several *Drosophila* species, the ab3 sensilla (characterised by its larger amplitude “A” neuron) is known to exhibit within-species and interspecies variation in odorant responses^33-35^. This sensillum type plays a major role in ecological adaptations to feeding ^36,37^, and in egg-laying decisions ^38^.

Only *D. novamexicana* possessed a *Nov*ab5 sensillum housing an “A” neuron responding to benzyl butyrate and a “B” neuron excited by isopropyl benzoate. Out of the 200 basiconic sensilla recorded from in the *D. ezoana* and *D. virilis* funiculus, no sister types of this sensillum were identified. *D. novamexicana* did not display a sensillum type equivalent to *Ezo*ab5 and *Vir*ab5. In *D. ezoana* and *D. novamexicana*, on the other hand, we could not find the sister type of the *Vir*ab11 sensillum. Several studies (reviewed in Anholt^39^) have shown that the ecological adaptation of drosophilid flies to specific environments is accompanied by rapid evolutionary changes in their olfactory system. Absence of specific sensillum types has e.g. been noticed in *D. sechellia*, where Stensmyr et al.^35^ observed the loss of the ab2 sensillum type paralleled by an over-representation of the ab3 type. This change has been hypothesized to be driven by adaptation to the food source of *D. sechellia*, the noni fruit. The loss of specific sensillum types in *D. ezoana* and *D. novamexicana* could also be the result of adaptations to their lifestyle, which is restricted to arctic and desert regions, respectively. Still, all three species under study here find their food in tree sap from a restricted spectrum of tree species, which would provide similar selection pressures in shaping the sensillum arsenal.

A broad behavioral two-choice experiment revealed the preferences of the three species when presented with 20 key ligands from the SSR experiments at a fixed testing temperature of 23°C. Some of the tested compounds elicited differential behavioural responses in the species. For instance, acetoin and ethyl-3-hydroxybutyrate elicited clear attraction in *D. virilis* but no response in *D. ezoana* and *D. novamexicana*. Methyl benzoate and 6-methyl-5-hepten-2-one attracted significantly fewer flies than the control in *D. novamexicana* and *virilis*, while it elicited a neutral response in *D. ezoana*. Such differences have also been observed among other fruitfly species. Dweck et al.^40^ found that 4-ethylguaiacol and methyleugenol derived from yeast fermentation induce strong attraction in *D. melanogaster* but elicit no response in *D. suzukii*. Similarly, the avoidance behaviour triggered by CO2 in *D. melanogaster* is not conserved in *D. suzukii* ^41^. The behavioural differences observed between the present three species might represent a taxon-specific adaptation to their respective environement. Such ecological adaptations are often accompanied by functional changes in some olfactory circuits ^42,43^, leading to situations where some odours elicit strong physiological responses but weak behavioural responses or vice versa ^44^.

In contrast, we found that guaiacol, methyl salicylate and isopropyl benzoate elicited conserved avoidance, attraction and neutral response, respectively, in *D. ezoana*, *D. novamexicana* and *D. virilis*. This suggests that the olfactory circuits responsible for the detection of these compounds are possibly conserved in the three species. Insects possess several olfactory circuits that are highly conserved, allowing them to accomplish complex behaviors, such as courtship, feeding, oviposition avoidance of enemies or toxic microorganisms^45^ . An interesting example to such a circuit is the geosmin dedicated olfactory circuitry^46^. This odour, emitted by molds, activates a specific, narrowly tuned olfactory receptor expressed in the ab4B neuron (Or56a) in the majority of drosophilid species (including the three studied here). In *D. melanogaster* specifically, the stimulation of the ab4B OSNs by geosmin triggers activity in a single glomerulus (DA2) and inhibits feeding, attraction and oviposition ^46^. However, if guaiacol (conserved repellent in our study) acts as antifeedant, attraction inhibitor, or oviposition deterrent like geosmin does to *D. melanogaster* still remains to be elucidated. Nonetheless, in another Diptera system, guaiacol (one of the waterbuck body odours) was found among the main odours composing the formulation (Waterbuck Repellent Blend (WRB)) that repels the tsetse fly species *Glossina fuscipes fuscipes*^47^. Further, methyl salicylate, identified as an attractant for the three species tested here, is an herbivore-induced plant volatile (HIPV) known to attract herbivore predators and parasitoids ^48^. In *D. melanogaster*, this odour exhibits a neutral valence when tested in a trap assay ^49^. Yellow traps baited with methyl salicylate significantly enhances catches of the dance fly, *Rhamphomyia gibba* (Diptera, Empididae) ^50^.

The temperature-modulated behavioural response assays revealed that, when tested at the extreme temperatures of 10 and 35°C, a large proportion of flies in all three species remained passive outside the traps, not making a choice to enter, indicating that these two extreme temperatures significantly impacted the ability to orientate and move. Recently, Ito and Awasaki^51^ also showed that the locomotor activity in 11 *Drosophila* flies (including *D. virilis*) matched the temperatures they frequently encounter in their habitats. As the species tested in our study were less likely to be active at 10°C or 35°C in their environment and showed both impaired activity in the choice assay at these temperatures (less than 50% made a choice in the two choice bioassay), and a clear avoidance of the same in our thermal preference assay, we decided to remove these data from further consideration.

In the remaining data, where flies reared at 18 and 25°C were tested for their preference at 15, 20, 25 and 30C, we observed some temperature-dependent effects. Below we discuss these for each of the compounds tested.

### Guaiacol (repulsive)

For this compound of general negative behavioral valence we observed temperature-dependent modulation of responses. In the cold- and warm-loving species the negative response was abolished or even reversed when in suboptimal temperature environments. In *D. ezoana*, repulsion disappeared at higher temperatures and was even reversed to attraction in flies reared at 25C. In *D. novamexicana*, a species adapted to warmer climates, we observed the opposite temperature response, with lower testing temperatures abolishing the negative response to guaiacol. In the temperate *D. virilis*, the response remained largely unaffected over the temperatures tested. Taken together, these results indicate that the ambient testing temperature had a clear impact on the behavior towards guaiacol, while the developmental temperature had little effect on this behavior. At less permissive temperatures the behavior vanished or was even reversed. Failing to respond to signals of danger might significantly reduce the survival of these flies, as this behavior often serves to escape pathogens, parasitoids and predators. For instance, Hangartner et al.^52^ found that *D. melanogaster* exposed to extreme heat had a lower survival rate when faced with predation. These authors postulated that extreme temperatures affect the flies’ physiological performance and impair their ability to detect and escape predators.

### Methyl salicylate (attractive)

Development at suboptimal temperatures significantly impacted the attraction behaviour in all three *Drosophila* species. Attraction also appeared only in a very narrow temperature window suggesting that exposing the preimaginal or adult stages of flies to uncommon temperatures can modulate their odour-guided attraction. Several studies outside the field of olfaction have pointed in a similar direction. For example, it was found that at low temperatures, the feeding preference for yeast in *D. melanogaster* was strongly reduced ^53^. In *D. suzikii*, the oviposition activity on blueberries decreased continuously above 28 °C and below 15°C ^54^. Our results also show the importance of the temperature in the environment where the behavior is performed. All three species displayed attraction to methyl salicylate in a 5-10 degree window, which seems lower in the cold loving species as compared to the temperate and warm-living ones.

### Isopropyl benzoate (neutral)

Very little impact of rearing and testing temperatures was observed when testing the compound that elicited no choice at 23C. In *D. ezoana* it remained neutral through all tested temperatures and under both rearing regimes. In the other two species a few tests showed weak attraction or repulsion but did not reveal any clear patterns.

In summary, our study showed that drosophilid flies choose locations with a temperature matching their natural habitats. We also found that temperature modulation can affect odor-guided behavior in three drosophilid species of different geographical and climatic origins. By recording olfactory responses we established relevant stimuli of differing behavioral significance and further demonstrated that the innate thermal preference of the three drosophilids to some extent also dictates their odor-mediated behaviors to ecologically relevant odors. The innate valence of such compounds can thus change at suboptimal temperatures outside of the innate thermal range. The mechanisms underpinning the changes observed might reside at different levels, from sensory detection to muscle action. In the future, it will be worth investigating the cellular and molecular mechanisms underlying the plastic character of olfactory mediated behavior of drosophilid flies when exposed to different temperatures.

## Acknowledgements

We thank Silke Trautheim for helping with the ordering of the three Drosophilid species used in these study. We thank also Daniel Veit for conceiving the thermal gradient apparatus used to test the temperature preference in the three drosophilid flies

## Author contributions

S.B.S.S, M.K, and B.S.H. designed the research plan. S.B.B.S performed all the experiments, analysed the data and prepared the figures. S.B.S.S, VPM, M.K, and B.S.H. discussed the results and wrote the manuscript.

## Data availability

The raw data generated and analysed in the current study are available from the corresponding author on reasonable request.

## Supplementary figures

**Supplementary figure 1.**
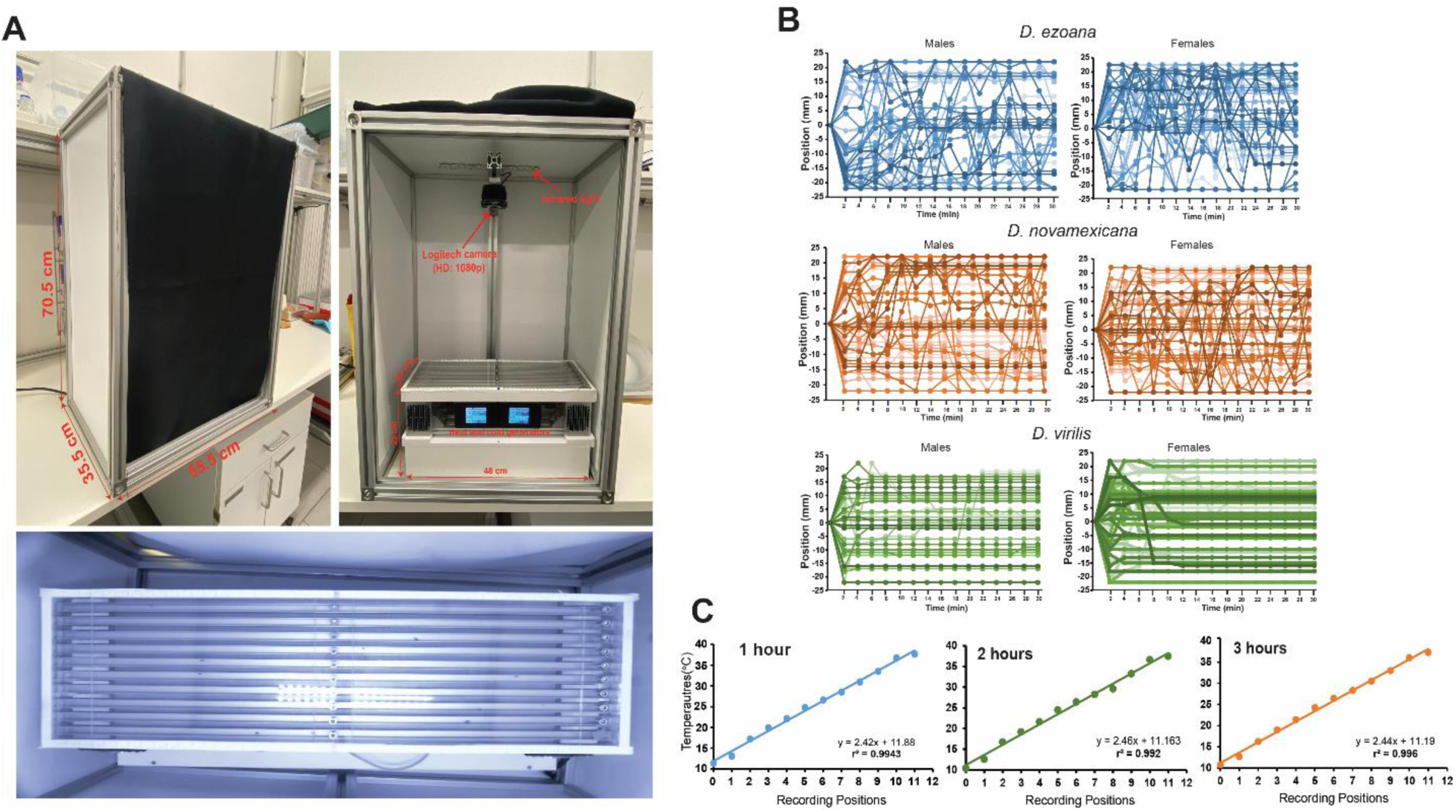
**(A)** Experimental setup for testing temperature preference in the three drosophilid flies. **(B)** Tracking lines showing the longitudinal position (recorded every 2 min) of *D. ezoana*, *D. novamexicana*, and *D. virilis* within the aluminium channels in absence of a thermal gradient. **(C)** Line graphs depicting the increase of temperature in the aluminium channels (from the cold region to the hot region) 1, 2 and 3 hours after turning on the heat and cold generators. Each dot on the graphs represents the temperature of a specific region of the aluminium channels.

**Supplementary figure 2.**
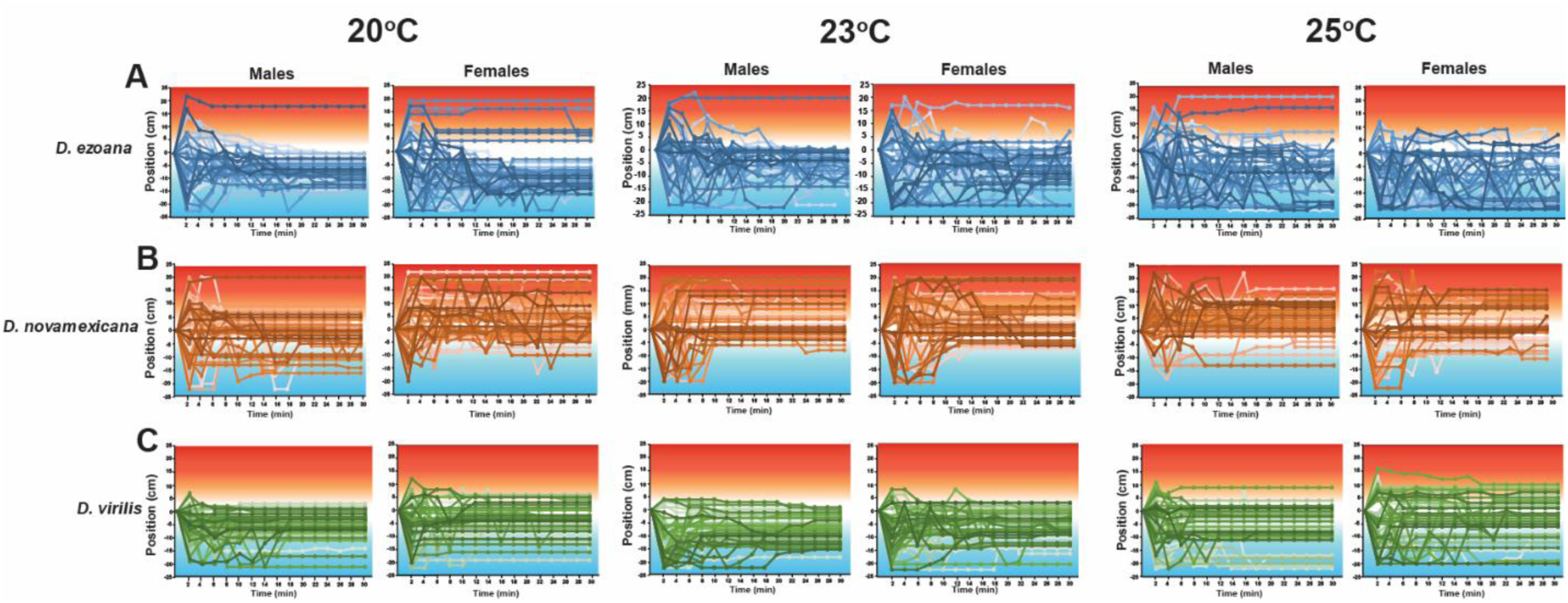
Tracking lines showing the longitudinal position (recorded every 2 min) of *D. ezoana* (**A**), *D. novamexicana* (**B**), and *D. virilis* (**C**) individuals reared at 20°C (left), 23°C (middle) and 25 °C right) and placed within the aluminium channels in presence of the thermal gradient.

**Supplementary figure 3.**
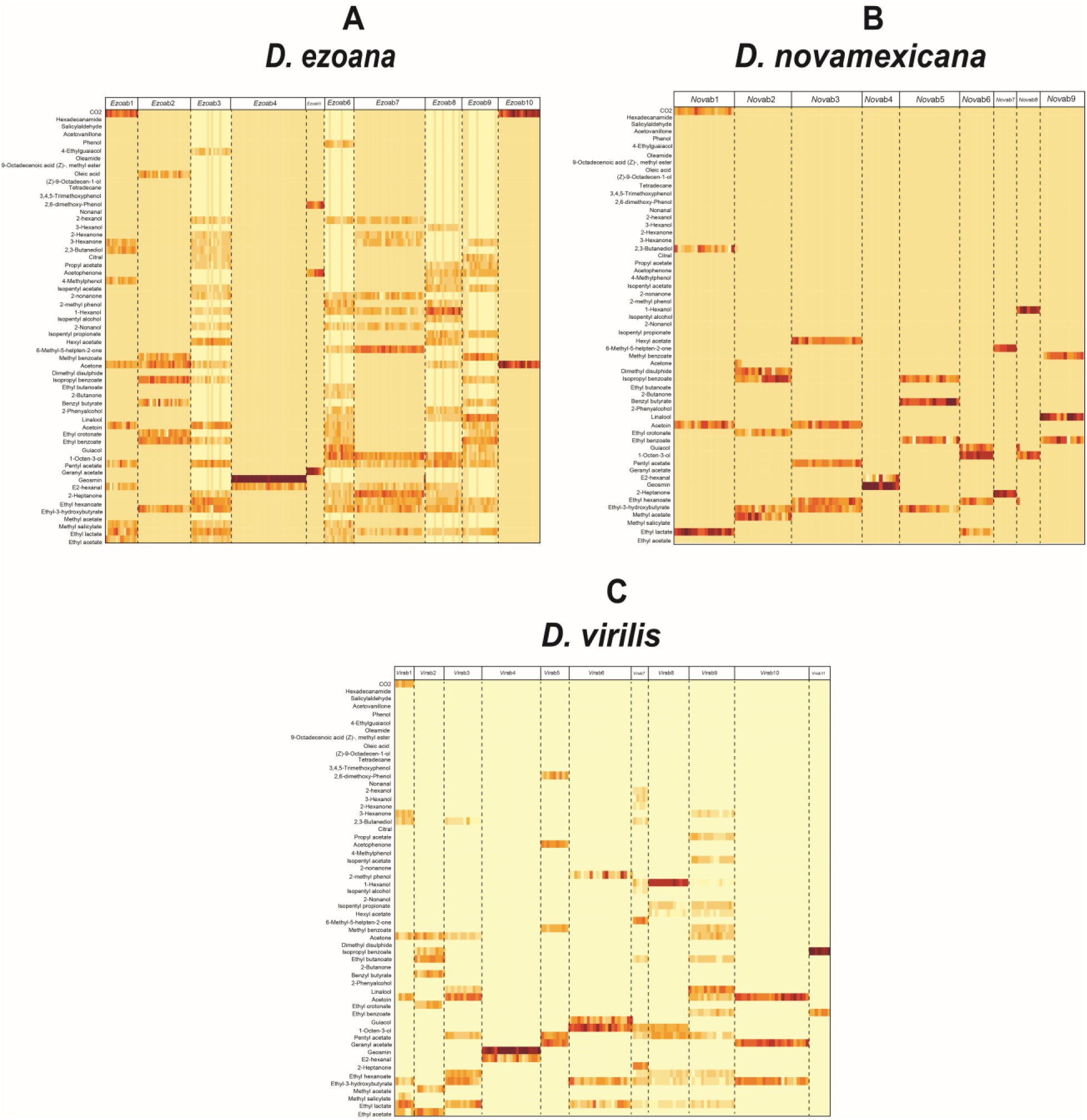
Functional classification of the antennal basiconic sensilla in the three *Drosophila* species. Heat maps show the response profiles of the 200 individual basiconics stimulated with the 57 odorant panel in (**A**) *D. ezoana*, (**B**) *D. novamexicana* and (**C**) *D. virilis*. Red indicates a maximum response; dark-orange a medium response; and yellow no response.

**Supplementary figure 4.**
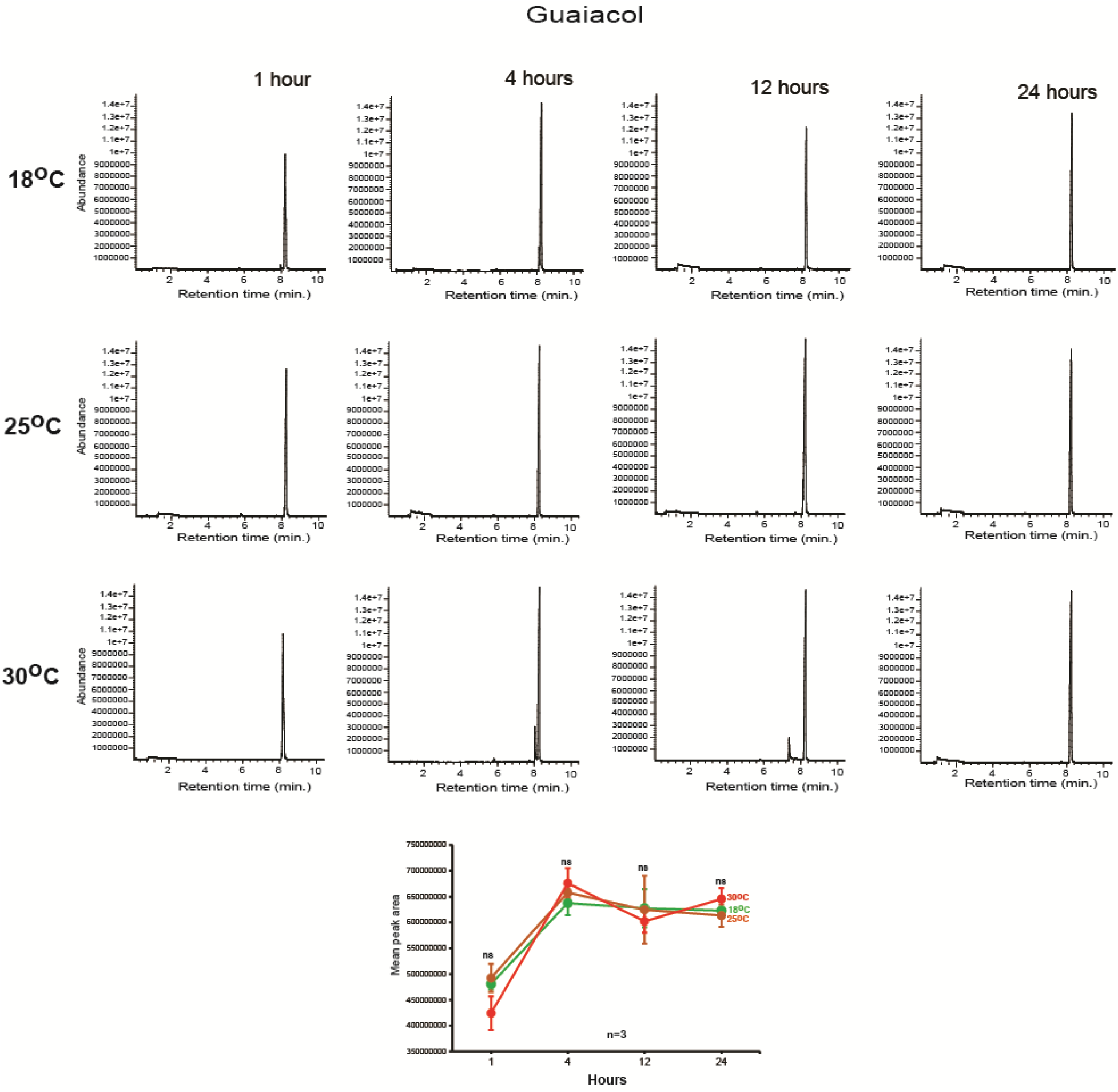
Release rate of Guaiacol in function of temperature and time

**Supplementary figure 5.**
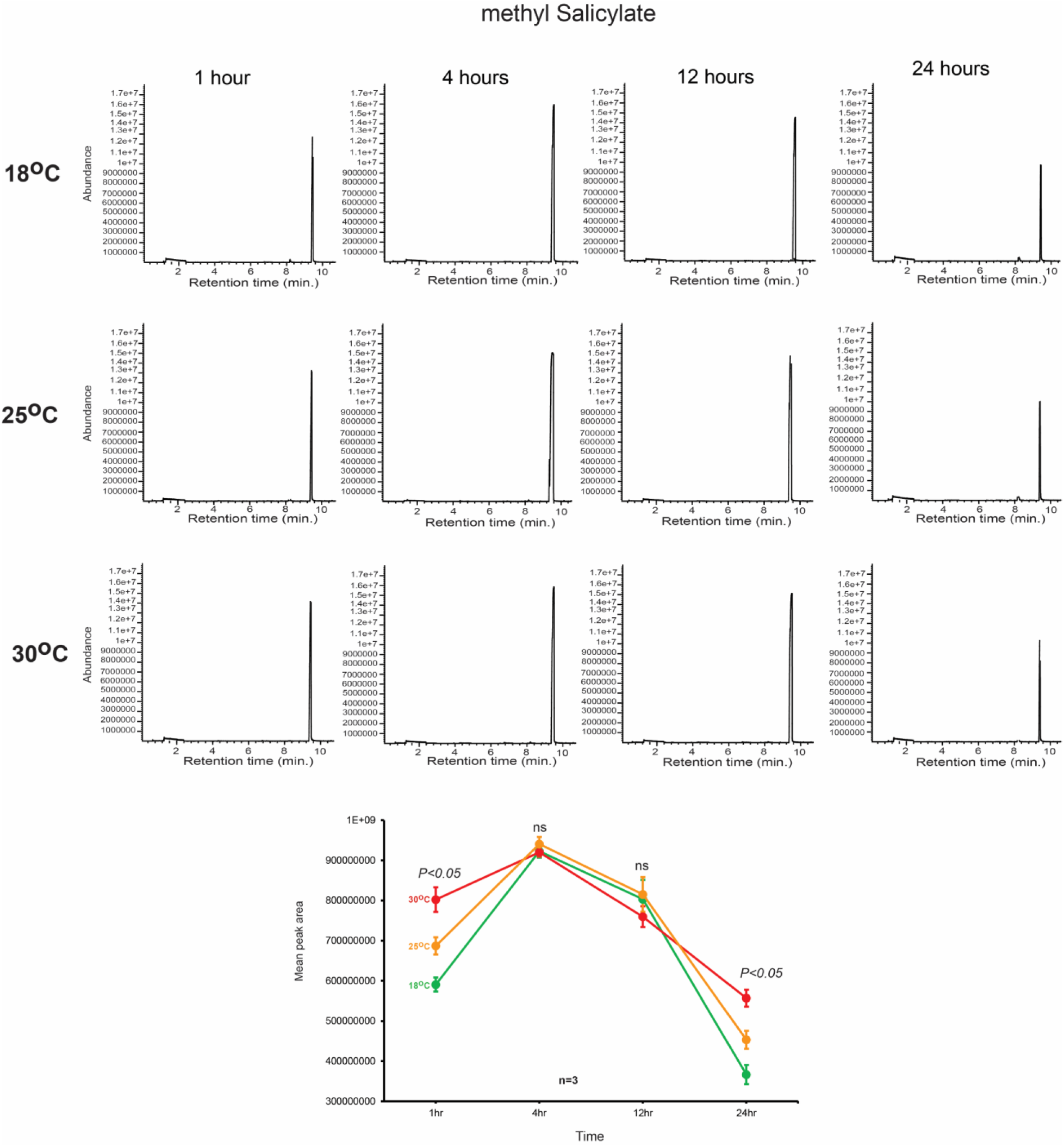
Release rate of methyl salicylate in function of temperature and time

